# Exposure to parasitic protists and helminths changes the intestinal community structure of bacterial microbiota but not of eukaryotes in a cohort of mother-child binomial from a semi-rural setting in Mexico

**DOI:** 10.1101/717165

**Authors:** Oswaldo Partida-Rodriguez, Miriam Nieves-Ramirez, Isabelle Laforest-Lapointe, Eric Brown, Laura Parfrey, Lisa Reynolds, Alicia Valadez-Salazar, Lisa Thorson, Patricia Morán, Enrique Gonzalez, Edgar Rascon, Ulises Magaña, Eric Hernandez, Liliana Rojas-V, Javier Torres, Marie Claire Arrieta, Cecilia Ximenez, Brett Finlay

**Affiliations:** Laboratorio de Inmunología del Departamento de Medicina Experimental, UNAM, Mexico City, Mexico; Michael Smith Laboratories, Department of Microbiology & Immunology, University of British Columbia, Vancouver, British Columbia, Canada; Department of Physiology & Pharmacology, University of Calgary, Calgary, Alberta, Canada; Department of Pediatrics, University of Calgary, Calgary, Alberta, Canada; Department of Zoology, University of British Columbia, Vancouver, British Columbia, Canada; Department of Botany, University of British Columbia, Vancouver, British Columbia, Canada; Unidad de Investigación en Enfermedades Infecciosas, UMAE Pediatria, IMSS, Mexico City, Mexico; Department of Biochemistry and Molecular Biology, University of British Columbia, Vancouver, British Columbia, Canada; Department of Microbiology and Immunology, University of British Columbia, Vancouver, British Columbia, Canada

**Keywords:** Parasites, Eukaryotes, Protists, Helmints, Microbiota, Bacteria, 16S sequencing, 18S sequencing, Children, Mexico

## Abstract

Around 3.5 billion people are colonized by intestinal parasites worldwide. Intestinal parasitic eukaryotes interact not only with the host, but also with the intestinal microbiota. In this work, we studied the relationship between the presence of multiple enteric parasites and the community structure of the bacterial and eukaryote intestinal microbiota in an asymptomatic cohort of mother-child binomials from a semi-rural community in Mexico. The intestinal parasites identified were *Blastocystis hominis, Entamoeba histolytica/dispar, Endolimax nana, Chilomastix mesnili, Iodamoeba butshlii, Entamoeba coli, Hymenolepis nana* and *Ascaris lumbricoides.* We sequenced bacterial 16S rDNA and eukaryotic 18S rDNA in fecal samples of 46 mothers and their respective children, with ages ranging from two to twenty months. Although we did not find significant alpha-diversity changes, we found a significant effect of parasite exposure on bacterial beta-diversity, which explained between 5.2% and 15.0% of the variation of the bacterial community structure. Additionally, exposure to parasites was associated with significant changes in relative abundances of bacterial taxa, characterized by increases in the Clostridia and decreased Actinobacteria and Bacteroidia abundances. There were no significant changes of intestinal microeukaryote abundances associated with parasite exposure. However, we found several significant positive correlations between intestinal bacteria and eukaryotes, including co-occurrence of the fungi *Candida tropicalis* with *Bacteroides* and Actinomyces, and Saccharomycetales with *Bifidobacterium* and *Prevotella copri.* These bacterial community structure changes associated with parasite exposure imply effects on microbial metabolic routes, host nutrient uptake abilities and intestinal immunity regulation in host-parasite interactions.

**IMPORTANCE:** The impact of intestinal eukaryotes on the prokaryotic microbiome composition of asymptomatic carriers has not been extensively explored, especially in children and in hosts with multiple parasites. In this work, we studied the relationship between protist and helminth parasite colonization and intestinal microbiota structure in an asymptomatic population of mother-child binomials from the semi-rural community of Morelos in Mexico. We found that the presence of parasitic eukaryotes correlated with changes in the bacterial community structure in the intestinal microbiota in an age-dependent way. This was characterized by an increase of the relative abundance of the class Clostridia and the decrease of Actinobacteria and Bacteroidia. While there were no significant associations between the presence of parasites and microeukaryote community structure, we observed strong positive correlations between bacterial and eukaryote taxa, identifying novel relationships between prokaryotes and fungi, and reflecting the diet of the human population studied.

## INTRODUCTION

Bacteria, viruses, archaea, fungi and protists inhabiting the mucosal surfaces of the human body have coevolved with the human intestine for millions of years and have broad effects on host metabolism and immunity. Under natural conditions, the diverse intestinal bacterial community shares their habitat with a dynamic community of eukaryotes, many of which are well-known parasites. Their interaction can affect the success of parasite colonization, influencing its outcome along the entire parasitism-mutualism spectrum (1), affecting multiple physiological processes, including metabolism, and normal development and function of the mucosal and systemic immune system (2–10).

The influence of intestinal parasites in resident bacterial and eukaryotic community structures has not been fully addressed. Some of the previous studies have been performed in experimental models of disease (10). Even though parasites colonize millions of people around the world, very frequently the damage caused by parasitic colonization is either controlled by the host or by both parasite and host, generating an asymptomatic colonization. Thus, the study of how parasites influence intestinal microbiota in asymptomatic individuals is an important and a relevant topic, especially given the large effect the microbiome can have on the host.

Little is known about the eukaryotic microbiome or eukaryome, and virtually nothing is known about its community structure, particularly in the case of protist colonisations. Morton et al. (5) found a strong association of the presence of *Entamoeba* with a higher frequency of Firmicutes and a lower frequency of *Bacteroidetes* in *E. histolytica-positive* samples. Nieves-Ramirez *et al.* (11), working in the same semi-rural Mexican population that is used in this study, found that asymptomatic colonization with the protist *Blastocystis* in adults was strongly associated with an increase in alpha- and beta-diversities of bacteria, and with more discrete changes in the microbial eukaryome.

It has already been widely documented that the initial development of intestinal microbiota in children has a profound effect on adult intestinal health and disease (12–16). The first 1,000 days of life play an important role in determining the phylogenetic structure of adult human gut microbiota (16) and early postnatal exposures to challenges, such as parasite exposure, could directly affect development of the gut microbiota structure. Given our lack of knowledge on the role of eukaryotes in the establishment of the early life microbiota, we aimed to study this in a population of 46 mother-child asymptomatic binomials from a semi-rural Mexican population with high levels of intestinal parasite exposure.

In this work, a comprehensive microbiome assessment was performed using 16S ribosomal DNA (rDNA) and 18S rDNA Illumina sequencing analysis for the characterization of bacteria and eukaryotes in the fecal microbiome Alpha and beta diversity, and bacterial relative abundance were determined and correlations between the parasite-positive and negative data sets revealed interesting associations between parasite colonization and distinct microbiome patterns in the intestine of children under two years of age.

## RESULTS

### Demographics

A Mother-Child (M-C) binomials cohort, which included parasite-positive individuals (23 M and 11 C), parasite-negative individuals (23 M and 23 C) and 12 children exposed to parasites (negative children with positive mothers) was studied (Table 1). The ages of the children ranged from two to twenty months (average of 11.04 months). Children were breast fed for an average of 11.2 months (2-20 months), with a significantly lower breast-feeding length in the parasite-positive (10 months) when compared to parasite-negative ones (12.43 months, p=0.0406). Twelve children were delivered by C-section and 34 through vaginal birth. All individuals were within a healthy weight range and did not show any stunting or wasting. Mother’s age average was 27.4 years old with a range of (18-47 years old). The mothers of parasite-positive children were significantly younger (25.3 years old) in comparison with the mothers of parasite-negative children (29.5 years old, p=0.0196). None of the participants reported antibiotic use, gastrointestinal symptoms (according to Rome III questionnaire), nor inflammatory signs and symptoms (according to medical examination) in the 6 months prior to sampling.

**TABLE 1.** Demographic data of the children in this study

|  | Parasite-positive<br>n=23 | Parasite-negative<br>n=23 | p value* |
| --- | --- | --- | --- |
| Age mean (months) | 11.09 | 11.00 | 0.9713 |
| Age mean of weaning<br>(months) | 10.00 | 12.43 | 0.0406 |
| Sex (female/male) | 12/11 | 8/15 | 0.2342 |
| Mode of delivery<br>(vaginal/C-section) | 17/6 | 17/6 | >0.9999 |
| Mothers age mean<br>(years) | 25.35 | 29.50 | 0.0196 |

\*Statistical analysis was made using Chi-square test for categorical data and t-test for numerical data.

### Parasites found in mother-child binomials

Intestinal parasites found either by microscopy or qPCR in feces of mothers and children included *Blastocystis hominis, Entamoeba coli, Entamoeba histolytica/dispar, Endolimax nana, Iodamoeba butshlii, Giardia duodenalis, Chilomastix mesnili, Hymenolepis nana* and *Ascaris lumbricoides* (Fig. 1). The most frequently found parasite was *B. hominis,* present in 32.6% of positive mothers and in 8.7% of children, followed by *E. coli* (M=23.9% and C= 8.7%). In the mothers, it was also common to find *E. histolytica/dispar* (10.8%), *E. nana* (10.8%), and *H. nana* (6.5%). In the children, *A. lumbricoides* was present in 8.7% of the positive samples. We also found several co-colonisations by two or more intestinal parasites particularly in the mothers, being *B. hominis-E. coli* the most frequently found (M=10.8% and C=2.2%). None of the parasite-positive individuals reported gastrointestinal symptoms.

**FIG. 1.**
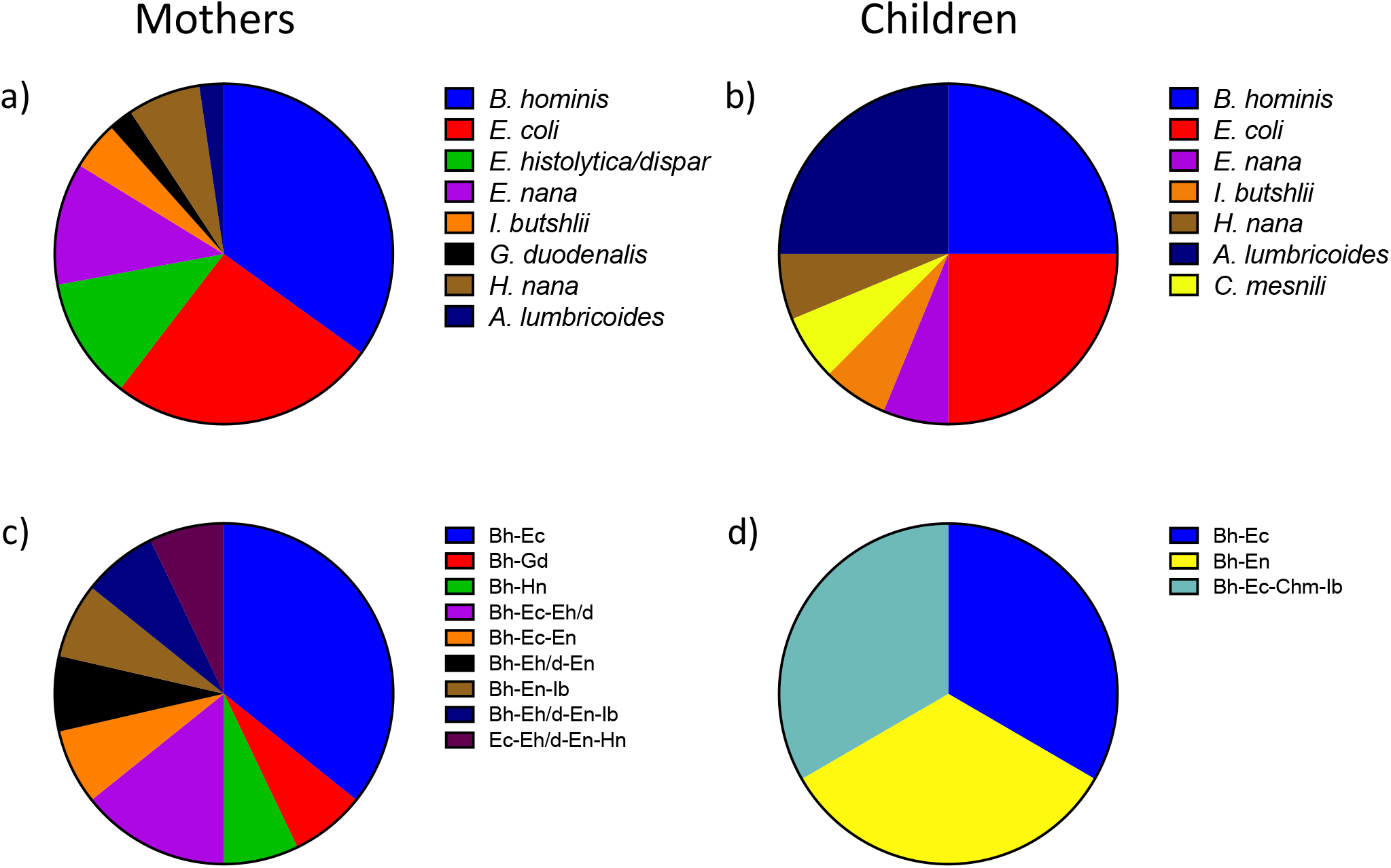
Proportions of parasites found in feces samples from mothers (a, c) and children (b, d) binomes. a) The intestinal parasites found in the mothers comprised *Blastocystis hominis* (Bh) [15 (32.6%)], *Entamoeba coli* (Ec) [11 (23.9%)], *Entamoeba histolytica/dispar* (Eh/d) [5 (10.8%)], *Endolimax nana* (En) [5 (10.8%)], *Iodamoeba butshlii* (Ib) [2 (4.3%)], *Giardia duodenalis* (Gd) [1 (2.2%)], *Chilomastix mesnili* (Chm) [0 (0.0%)], *Hymenolepis nana* (Hn) [3 (6.5%)], and *Ascaris lumbricoides* (Al) [1 (2.2%)]. b) In children, the parasite frequencies found were Bh [4 (8.7%)], Ec [4 (8.7%)], Eh/d [0 (0.0%)], En [1 (2.2%)], Ib [1 (2.2%)], Gd [0 (0.0%)], Chm [1 (2.2%)], Hn [1 (2.2%)] and Al [4 (8.7%)]. c) Co-colonisations by two or more parasites found in the mothers included Bh-Ec [5 (10.8%)], Bh-Gd [1 (2.2%)], Bh-Hn [1 (2.2%)], Bh-Ec-Eh/d [2 (4.3%)], Bh-Ec-En [1 (2.2%)], Bh-Eh/d-En [1 (2.2%)], Bh-En-Ib [1 (2.2%)], Bh-Eh/d-En-Ib [1 (2.2%)] and Ec-Eh/d-En-Hn [1 (2.2%)]. d) Co-colonisations found in children were Bh-Ec [1 (2.2%)], Bh-En [1 (2.2%)], and Bh-Ec-Chm-Ib [1 (2.2%)].

### Bacterial and eukaryotic diversity in individuals colonized by protists

#### Binome identity

We determined the fecal bacterial and eukaryotic composition of the 46 binomials. In order to evaluate if the mother’s intestinal microbiota composition could be a variation driver of the intestinal microbiota in children, we evaluated the binome identity on the beta-diversity by permutational multivariate analysis of variance (PERMANOVA) (11) on Bray-Curtis distances of bacteria and eukaryotes among mothers and their children. There were no observed significant effects of binome identity on the variation of either the bacterial or the eukaryote community structures in this group of individuals (Fig. S1).

##### Effects in children (less than five months old)

Due to the known differences in microbiome structure and diversity explained by temporal changes during the first year of human life (18), and that we observed a clear impact of age in the intestinal microbiota distribution due to age, we evaluated the effect of parasites in children under or over 1 year of age separately. For children under 1 year of age, which in this cohort were all under 5 months of age, we identified relationships between bacterial community structure and parasite exposure by conducting a PERMANOVA on the community matrix. We did not find any parasites among children younger than 5-months-old, but we considered them as parasite-exposed when their mothers tested positive for parasites. Based on Bray-Curtis dissimilarities (Fig. 2), we found that the community structure of bacteria was significantly related to parasite exposure (PERMANOVA, p=0.003). According to the PERMANOVA analysis, parasite exposure explained 15% of the variation in bacteria beta-diversity (Fig. 2a), however, we did not find an effect of parasite exposure on eukaryote community structure (Fig. 2b). We also calculated the Shannon diversity index and Chao1 estimated richness of bacterial and eukaryotic communities in this group of children and we did not find any statistically significant differences between the individuals exposed vs non-exposed to parasites (Fig. 2c, d, e, and f).

**FIG. 2.**
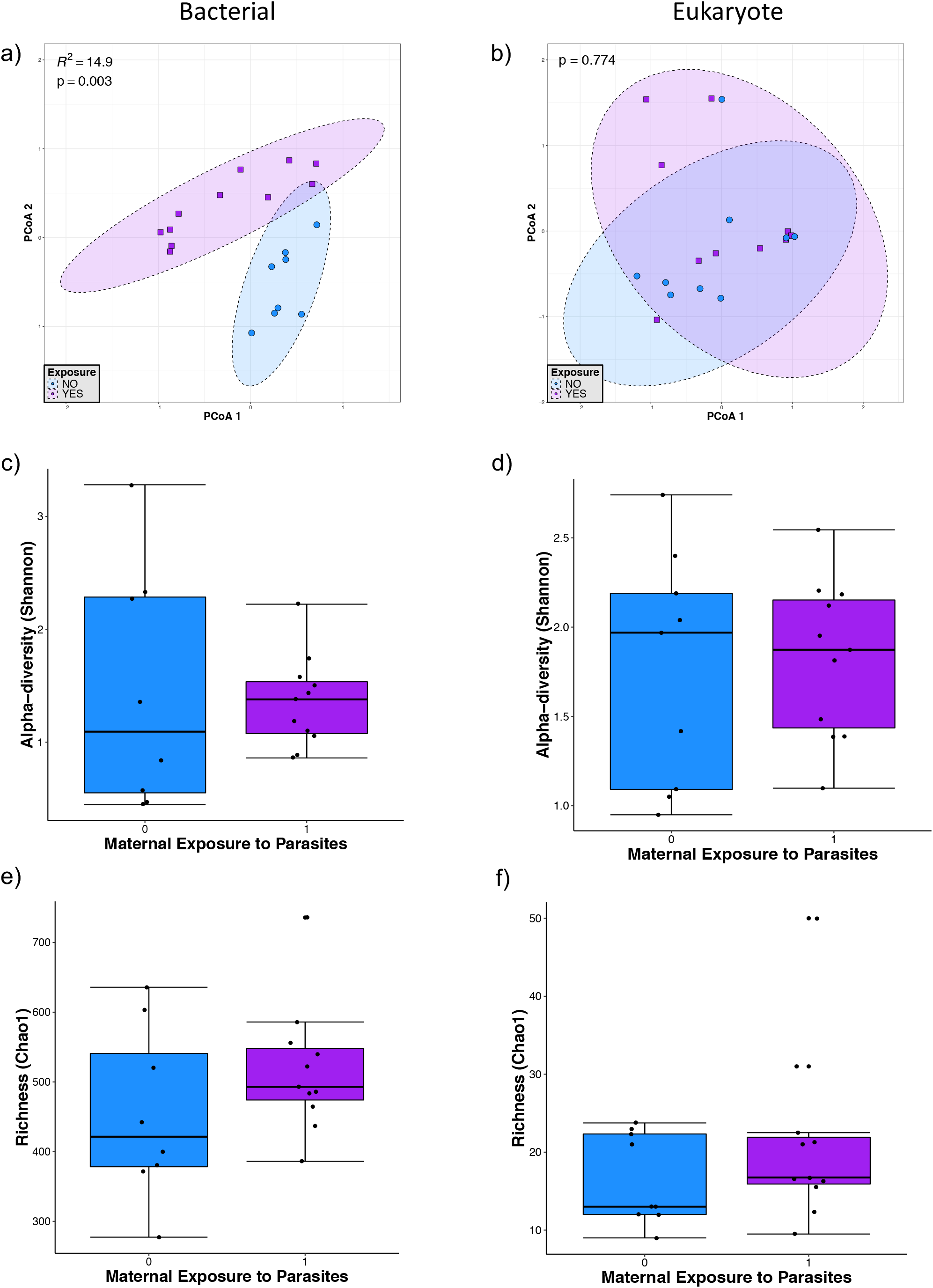
Microbial diversity in infants younger than 5 months of age. a) and b) Principal component analysis (PCoA) ordination of variation in beta-diversity of human gut bacterial (a) and eukaryote (b) communities based on Bray-Curtis dissimilarities. Color and shape represent maternal exposure to parasites (blue circles represent negative exposure and purple squares represent positive exposure). PERMANOVAs indicate that maternal exposure to parasites explain 15% (p=0.003) of the variation in infant bacterial community structure but is not a significant (p=0.774) driver of eukaryote community structure. c) and d) Shannon diversity of gut bacterial (c) and eukaryote (d) community structure. (e and f) Chao 1 estimated richness of gut bacterial (e) and eukaryote (f) community structure. No significant differences were detected by Mann-Whitney tests for alpha-diversity comparison between the parasite-positive and negative groups.

##### Children older than one year old

In children older than one year old, we found a significant effect of parasite presence on bacterial beta-diversity (Fig. 3a), which explained 5.2% of the variation present in this group. We also observed a significant effect of child age on both bacterial (p<0.001) (Fig. 3a) and eukaryote (p<0.001) (Fig. 3b) beta-diversities, explaining the variation found in 6.7% and 4.3%, respectively. Chao1 estimated richness in this group shown a significant increase of bacterial diversity (p=0.04) in the presence of intestinal parasites but not in the eukaryotic diversity (Fig. 3e). There were no statistically significant differences on Shannon diversity in bacteria and eukaryotes between parasite positive and negative children in this group of age (Fig. 3c and d), suggesting that changes in alpha-diversity are mainly due to effect on operational taxonomic unit (OTU) richness.

**FIG. 3.**
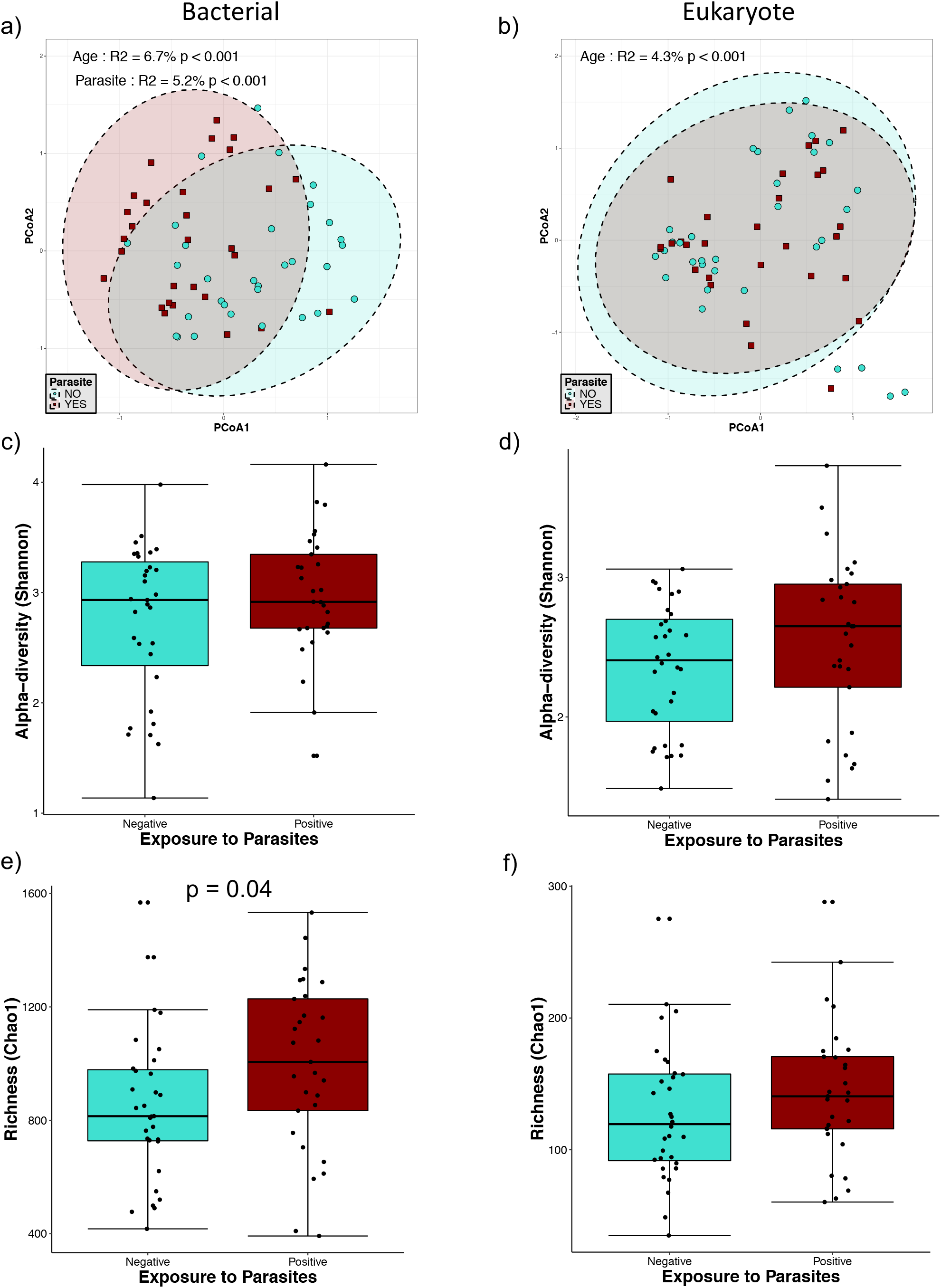
Microbial diversity in individuals older than 1yo. a) and b) Principal component analysis (PCoA) ordination of variation in beta-diversity of human gut bacterial (a) and eukaryote (b) communities based on Bray-Curtis dissimilarities. Color and shape represent maternal exposure to parasites (turquoise circles for negative and dark red squares for positive exposure). PERMANOVAs indicate that maternal exposure to parasites and age explain respectively 5.2% and 6.7% (p<0.001) of the variation bacterial community structure while age explains 4.3% (p<0.001) of the variation in eukaryote community structure. Ellipses represent confidence interval at 95%. c) and d) Shannon diversity of gut bacterial (c) and eukaryote (d) community structure, no significant differences were detected by Mann-Whitney tests for Shannon diversity between the parasite-positive and negative groups. e) and f) Chao 1 estimated richness of gut bacterial (e) and eukaryote (f) community structure; a significant difference was detected for bacterial community richness by Mann-Whitney tests for comparison between two groups.

##### Children from one to two years old

In older children, between one and two years old, we detected an effect of parasite presence on bacterial community structure (Fig. 4). The parasite exposure explained 8.7% of the bacterial variation whereas age explained 7.7% (Fig. 4a). Chao1 and Shannon diversity indices showed no changes in richness or evenness (Figure 4c and e). Although there was no significant effect detected on eukaryotic beta-diversity (Figure 4b) and on eukaryotic Chao1 richness (Fig. 4f), eukaryote alpha-diversity (Shannon index) showed a statistically significant increase in the group colonized by parasites (Fig. 4d).

**FIG. 4.**
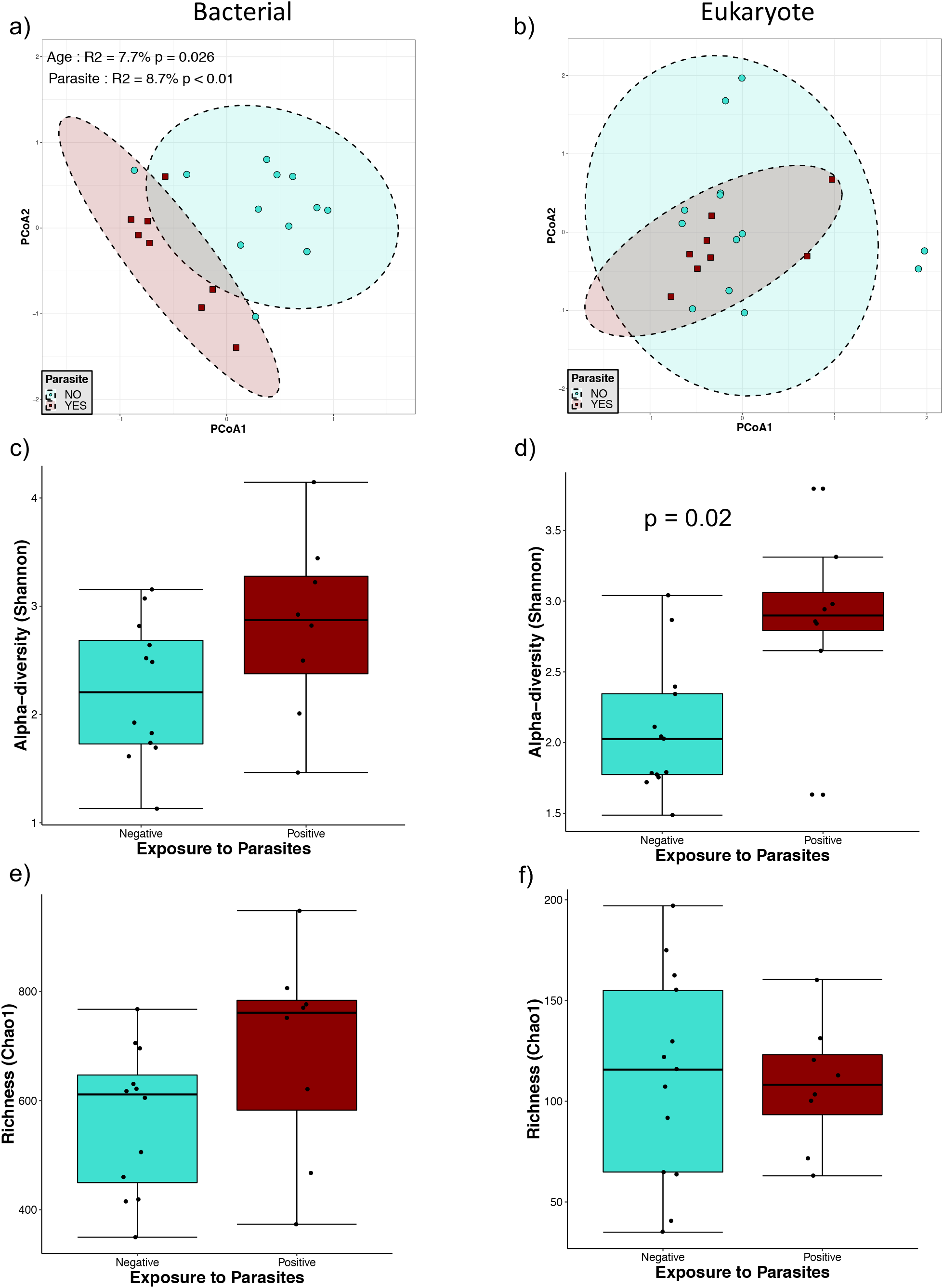
Microbial diversity in infants between 1yo and 2yo. a) and b) Principal component analysis (PCoA) ordination of variation in beta-diversity of human gut bacterial (a) and eukaryote (b) communities based on Bray-Curtis dissimilarities. Color and shape represent maternal exposure to parasites (turquoise circles for negative and dark red squares for positive exposure). PERMANOVAs indicate that exposure to parasites and age explain respectively 8.7% (p<0.01) and 7.7% (p=0.026) of the variation infant gut bacterial community structure. Ellipses represent confidence interval at 95%. No significant effects of age or exposure to parasite were detected for eukaryote community structure. c) and d) Shannon diversity of gut bacterial (c) and eukaryote (d) community structure, a significant difference was only detected for eukaryote community alpha-diversity by Mann-Whitney tests for comparison between the two groups. e) and f) Chao 1 estimated richness of gut bacterial (e) and eukaryote (f) community structure; no significant differences were detected by Mann-Whitney tests for Chao 1 estimated richness between the parasite-positive and negative groups.

In mothers, we also found a significant effect of parasite exposure on bacterial beta-diversity (Fig. 5a), explaining 5.6% of the variation of community structure, and no effect over the eukaryotic diversity (Fig. 5b). No statistically significant effects were observed in mothers exposed to parasite in bacterial and eukaryote community richness and alpha-diversity (Fig. 5c, d, e and f).

**FIG. 5.**
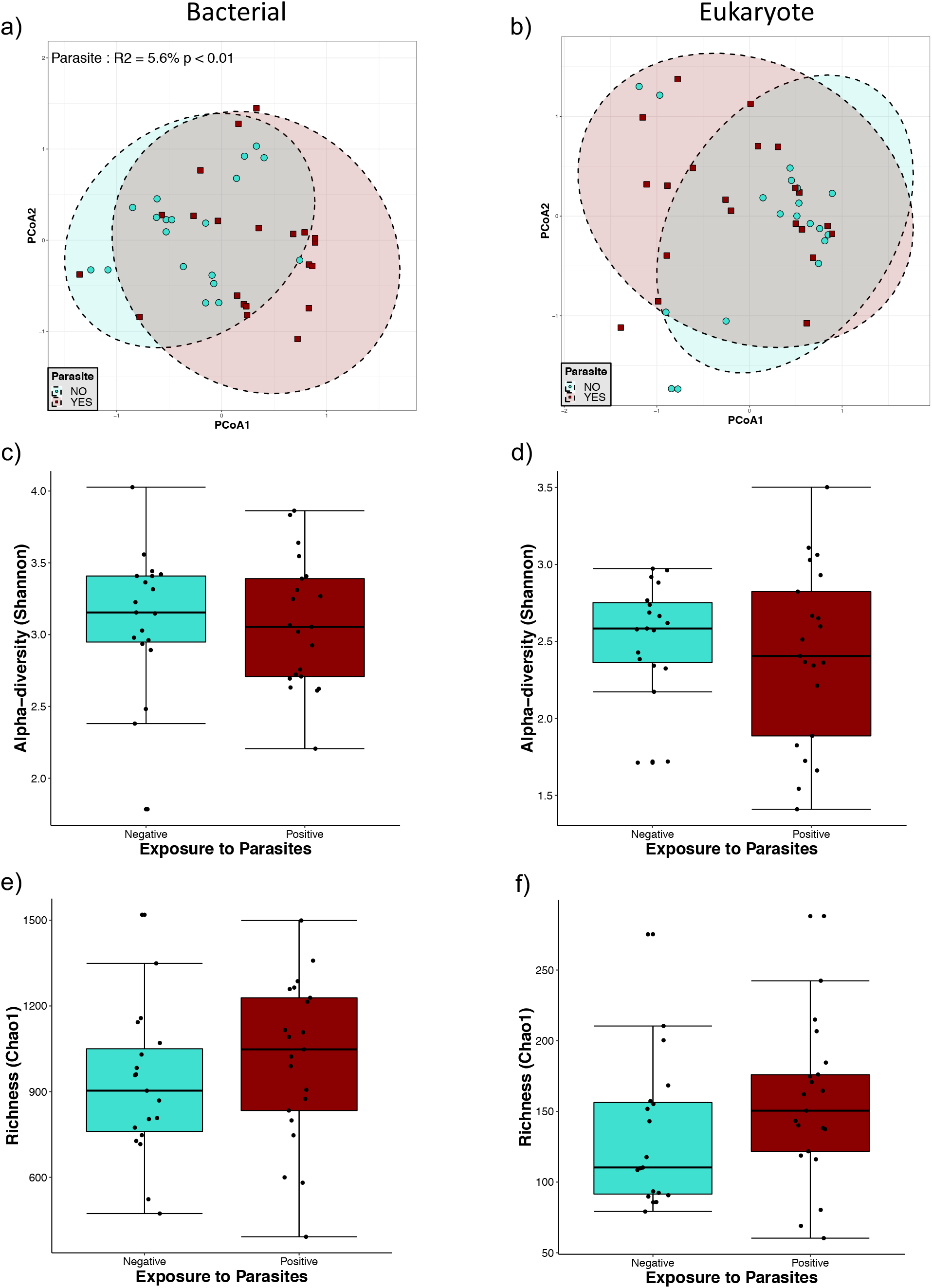
Microbial diversity in mothers. a) and b) Principal component analysis (PCoA) ordination of variation in beta-diversity of human gut bacterial (a) and eukaryote (b) communities based on Bray-Curtis dissimilarities. Color and shape represent maternal exposure to parasites (turquoise circles for negative and dark red squares for positive exposure). PERMANOVAs indicate that exposure to parasites explains 5.6% (p<0.01) of the variation in mother gut bacterial community structure. Ellipses represent confidence interval at 95%. No significant effects of age or exposure to parasite were detected for eukaryote community structure. c) and d) Shannon diversity of gut bacterial (c) and eukaryote (d) community structure. e) and f) Chao 1 estimated richness of gut bacterial (e) and eukaryote (f) community structure. No significant differences were detected by Mann-Whitney tests for richness and alpha-diversity between the parasite-positive and negative groups.

### Relative abundance of bacterial classes

Differential abundance analysis revealed changes in bacterial composition associated with parasite colonization and exposure (Fig. 6). In babies less than five months old exposed to parasites, the most abundant groups at the genus level were *Bifidobacterium* and *Bacteroides* (Fig. 6a). In this group of children, the genera *Pseudoramibacter, Eubacterium, Prevotella* and *Oscillospira* were reduced in their relative abundance compared to parasite exposure.

**FIG. 6.**
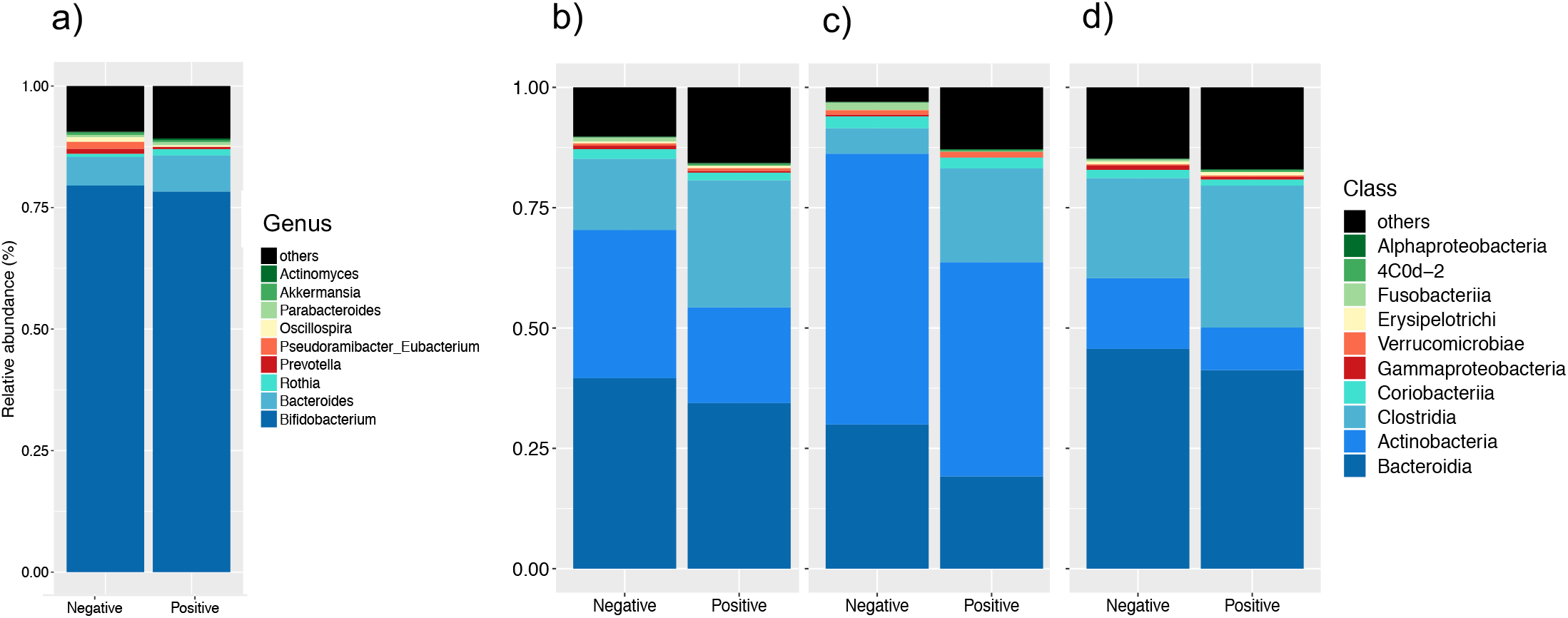
Relative abundance of gut bacterial composition of parasite-exposed individuals. a) Infants younger than 5 months of age bacterial composition at the genus level. Relative abundance at the class level depending on parasite colonization b) in individuals over 1yo; c) only in infants between 1yo and 2yo; and d) only in mothers.

In children older than one year, children from one to two years old and in the mothers, the most abundant classes were Bacteroidia, Actinobacteria, Clostridia and Coriobacteria, and the presence of parasites was consistently associated with an increase of the relative abundance of Clostridia and a decrease of Actinobacteria and Bacteroidia (Fig. 6b, c and d). There was also a decrease of Fusobacteria and Gammaproteobacteria percentages associated with the colonization of parasites in the children from one to two years old and in the mothers, respectively.

### Bacterial-Eukaryote correlations

In order to study correlations between the presence of bacterial and eukaryotic taxa, independently of the parasite exposure, we created heatmaps of biweight correlations between the top 50 bacterial taxon operational taxonomic units (OTUs) and the top 50 eukaryote OTUs in fecal samples. In babies less than five months of age, we found several statistically significant correlations between bacterial and eukaryote taxa (Fig. 7). We found positive correlations of bacteria with fungi: *Bacteroides* with *Candida tropicalis,* Eurotiales and *Hanseniaspora uvarum*; *Bacteroides fragilis* with Eurotiales and *Hanseniaspora uvarum*; Lachnospiraceae with Eurotiales and *Hanseniaspora uvarum*; Coriobacteraceae with Eurotiales and *Hanseniaspora uvarum*; *Oscillospira* with Nucletmycea, *Pichia kudinavzevii,* Eurotiales, and *Hanseniaspora uvarum;* Actinomyces with *Candida tropicalis*; and *Bifidobacterium* with Saccharomycetales. We also found positive correlations of *Bacteroides, Bacteroides fragilis,* Lachnospiraceae, Coriobacteraceae and *Oscillospira* with the protist *Heteromita.* In children older than one year old and in the mothers, there were less correlations between bacterial and eukaryote taxa (Fig. 8). We found positive correlations of *Prevotella copri* with *Saccharomyces*.

**FIG. 7.**
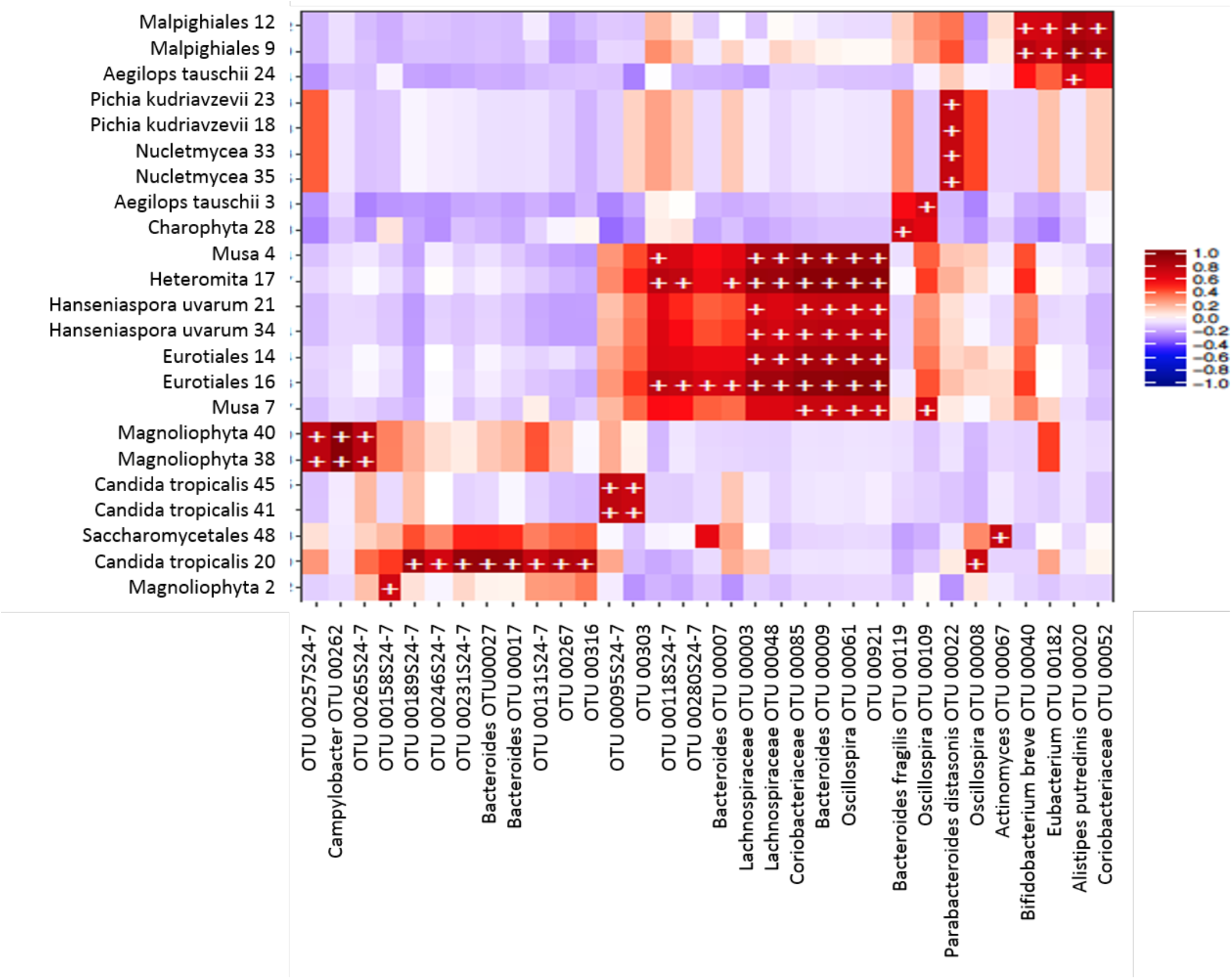
Heatmap of biweight correlations (Pearson) between the top 50 bacterial (x axis) and top 50 eukaryote taxon (*y* axis) OTUs in fecal samples of infants (~3-4 months-old). Colors denote positive (red) and negative (blue) correlation values. Significant correlations are denoted with a plus sign (*P* < 0.05; FDR).

**FIG. 8.**
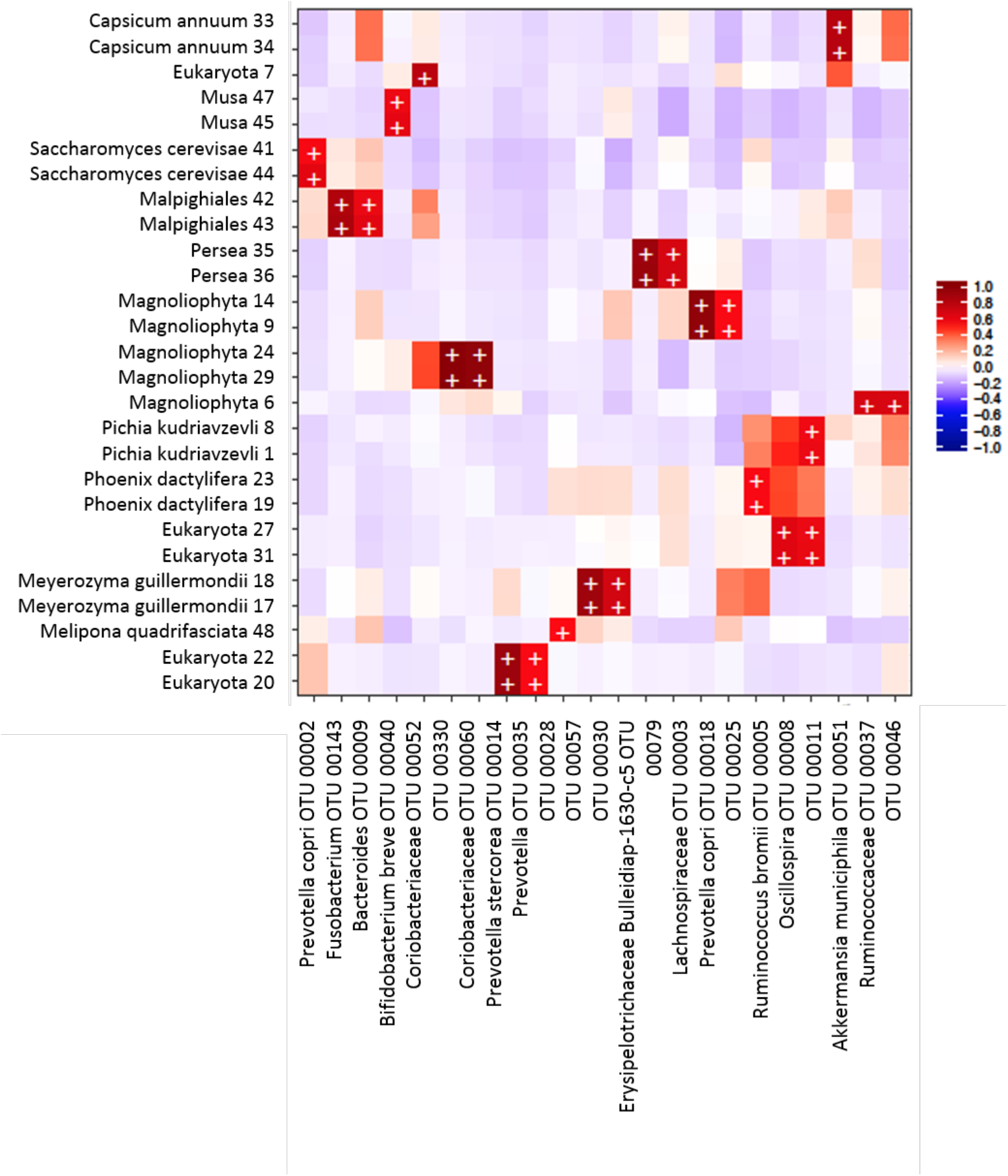
Heatmap of biweight correlations (Pearson) between the top 50 bacterial (x axis) and top 50 eukaryote taxon (*y* axis) OTUs in fecal samples of children (>1yo) and mothers. Colors denote positive (red) and negative (blue) correlation values. Significant correlations are denoted with a plus sign (P < 0.05; FDR).

## DISCUSSION

The most recent reports available estimate that 3.5 billion people are colonized by parasites around the globe (19, 20). Even though eukaryotes are in much lower abundance than bacteria, it has been demonstrated that mono-colonization of parasites is associated with intestinal microbiota composition changes (5, 11, 21–24). Colonization by parasitic eukaryotes usually does not follow a one host–one parasite model (25–29), and very few studies have assessed the intestinal microbiota composition when multiple parasites are present (30). In this project, we studied whether the exposure to intestinal parasites in an asymptomatic cohort of mother-child binomials from semi-rural community in Mexico is related to changes to the bacterial and eukaryotic intestinal microbiota. Asymptomatic cohorts are a crucial setting to study this research question, given that the inflammatory response in symptomatic parasitosis can confound the ecological effect of the presence of parasites in the human gut. Our study revealed important changes in bacterial intestinal microbiota relative abundances associated with the exposure to parasites, characterized by the increase of the abundance of the taxa Clostridia and the decrease of Actinobacteria and Bacteroidia, with no important changes in alpha-diversity indices.

We found nine different parasites in the binomials studied, predominated by the protists *Blastocystis hominis* (20.6%), *Entamoeba coli* (16.3%), *Endolimax nana* (6.5%) and *Entamoeba histolytica/dispar* (5.4%) and the two helmints *Ascaris lumbricoides* (5.4%) and *Hymenolepis nana* (4.3%). More than half of the exposed individuals (54.8%) were colonized by two or more different parasites, yet, all parasite-positive individuals in our cohort remain asymptomatic. Interestingly, when parasite positive individuals were compared with parasite-negative ones, we found that the parasite-positive children were breast-fed for a significantly shorter time and that the mothers of parasite-positive children were significantly younger. Even though we did not find parasite colonization in the children younger than 5-months-old, these infants are constantly exposed to the microbiome of their parasite-positive mothers. These exposures originate from birth, breastfeeding, and spending the majority of the time together (31). It remains unclear why infants at this age do not become colonized by parasites, but our data clearly indicates that even in the absence of colonization, there are distinct microbiome patterns associated with maternal colonization. These results suggest that bacterial microbiome differences originating from parasite colonization are inheritable and independent of colonization per se.

The mother’s microbiota has been determined as an important microbial source during early colonization of the infant gut in industrialized settings (13, 31–33), however, we found no binome identity effect either in bacterial or in eukaryote communities. This may be due to the fact that the majority of the abovementioned studies compared infant samples collected shortly after birth. The samples analyzed in this study were of older infants. However, it is also possible that the increased dissimilarity between mother-infant binomes in this study may reflect a bigger influence of other external environmental factors on infant microbiome structure in this semi-industrialized setting.

Our study detected an increase of the bacterial and eukaryotic richness and alpha-diversity due to parasite colonization only in children over 1 year. Previous reports have associated colonization with parasites with a higher intestinal bacterial diversity. The study by Morton et al. (5) in rural populations in Cameroon found that the presence of the protist *Entamoeba* was associated with a significant increase in alpha (intra-host) diversity. Furthermore, a recent study by our group (11) found a dramatic increase of the bacterial richness and alpha-diversity in people colonized by *Blastocystis* from the same community as the present study. The discrepancies between this study and previous ones, which did not include infants under 1 year, strongly suggest that the effect of parasite exposure on alpha-diversity is age-dependent, and may be attributable to the lower alpha-diversity in infant samples. This is further supported by a recent study in children from Colombia (30), in which no differences in bacteria alpha-diversity in children positive to parasites were reported either. However, we did not observe an increase in alpha-diversity in the parasite-positive mothers either, suggesting that other factors may be at play, like the common multiparasitic colonization in this settings that may lead to more complex interactions with the resident microbiota, which may only be detected in a much larger study population size.

Despite the discrepancies with other studies in relation to alpha-diversity, we found that the parasite exposure was significantly associated with the bacterial intestinal microbiota beta-diversity in both infants and mothers in this study yet not with the eukaryotic microbiota. In children younger than 5 months old, the parasite exposure explained 14.9% (p=0.003) of the variation in bacterial community structure, whereas in the older groups of infants and mothers parasite exposure explained less of an effect in beta-diversity. This suggests that while parasite exposure is an important factor shaping bacterial intestinal beta-diversity, its effects are more evident during the earlier stages of gut bacterial community establishment.

Exposure to parasites was also associated with changes in specific taxa, including an increase of bacterial class Clostridia and a decrease of class Bacteroidia. This result agrees with the study by Morton et al. (5), in which there was a strong correlation of *Entamoeba* with higher frequency of Firmicutes (particularly the Clostridia class) and a lower frequency of Bacteroidetes (mostly *Prevotella)* in *E. histolytica-positive* samples. In the study focused on *Blastocystis* (11), there was also a significant increase of the genera *Ruminococcus* and *Oscillospira* (members of the Clostridia class) and a reduction of the genus *Prevotella* (Bacteroidia class) in colonized individuals.

There are a several mechanisms possibly responsible for the observed changes in relative abundance. Direct parasite-bacteria interactions driven by competition for resources, predation or production of molecules may affect the fitness and survival or microorganisms involved (1). Protists are well-known bacterivores. *Entamoeba* and *Blastocystis* can graze on bacteria (34, 35), and their ability to feed on the bacteria is an important mechanism for top down control of bacterial communities due to their high feeding rates (36). It has also been recently reported that *Trichuris muris,* a nematode from the mouse gut, acquires its own intestinal microbiota from the mouse intestine, very likely through ingestion (37). Furthermore, the mechanisms by which bacteria avoid protists predation changes their ability to survive and can even promote the emergence of virulence and invasion (36, 38).

Protists may also influence bacterial community structure in the intestine through indirect interactions with bacteria. Some parasites, like *Entamoeba* and *Giardia* who have mucolytic enzymes (39, 40), and helminths like *Trichuris trichiura,* which stimulate mucin expression or express mucin-like molecules themselves (41–43), can alter the outer mucus layer changing the bacteria microenvironment and sources of nutrition for certain taxa. Additionally, the parasites may produce metabolites that could influence the regulation of the immune system, which helminths are well-known to do. *Trichuris muris* and *Heligmosomoides polygyrus bakeri* can induce the generation of Tregs (44, 45), changing the physical microenvironment by modifying the mucus and the antimicrobial peptides production, which might promote the outgrowth of specific species among the microbiota (46). Bacterial and eukaryotic taxa identified from our and other human studies should be studied in appropriate animal models to further determine mechanisms involved in these multi-kingdom interactions.

While exposure to parasites practically had no effect on the diversity and abundance of eukaryotes, correlation analysis detected several significant bacteria-eukaryotes positive associations. The strongest positive correlations were found between bacteria and several common fungi of the intestinal microbiota. Agonistic and antagonistic relationships have been described between intestinal fungi and bacteria (47–52). The common yeast *Candida albicans* suppressed regrowth of *Lactobacillus* and promoted the recovery of Bacteroidetes populations during antibiotic recovery (53). Our study detected co-occurrence of another *Candida* species, *Candida tropicalis,* with *Bacteroides.* Other co-occurrences between fungi and bacteria found were *Oscillospira* with Nucletmycea, *Pichia kudinavzevii,* Eurotiales, and *Hanseniaspora uvarum;* Actinomyces with *Candida tropicalis*; *Bifidobacterium* with Saccharomycetales, and *Prevotella copri* with *Saccharomyces.* While correlative in nature, our analysis may also reveal the results of bacteria-fungal interactions in the human gut.

Our work supports previous reports that presence of intestinal parasites is linked to strong bacterial microbiota community changes. By including mother-child binomes, our work further revealed that these effects occur even in the absence of direct colonization in infants of colonized mothers, strongly suggesting that the effect of parasite colonization on the microbiome may also lead to changes in the vertical transmission of bacterial taxa. This would imply that colonization by parasites may be a strong indirect factor in the inheritable features of the human gut microbiome. How these intestinal microbiota changes associated to parasites may modify the immune system and other aspects of metabolism remains to be elucidated.

## MATERIALS AND METHODS

### Study population, study design and ethical considerations

Xoxocotla is a semi-rural community of the State of Morelos, Mexico, located at 120 km south of Mexico City (longitude 99°19=W, latitude 18°3=N) in a spanning area of 29,917 km2 with a tropical climate (warm subhumid). Total population is 5,163 people whose main source of money income is agriculture and commerce. Sample collection was carried out between April 2011 and January 2013. In this cross-sectional study of cohorts, every volunteer mother was informed about the characteristics of the project, the objectives, and the advantages to participate, as well as the biological samples needed, the sampling procedures and possible complications that may arise. All the participant mothers signed a written informed consent letter for her child and herself prior to the sample collection. Afterwards, questionnaires to collect sociodemographic, socioeconomic and health data antecedents (Rome III for gastro-intestinal symptoms (54), nutrition, way of delivery, and antibiotic use about the 6 months prior sampling) were applied and all variables were recorded in a database. The recruited mothers were submitted to a stool microscopic analysis for detection of intestinal parasites for the construction of the parasitized and non-parasitized cohorts. Forty-six Mother-Child binomials were included in the study, and feces samples were collected in sterile plastic containers, immediately placed at 4°C for transport to the laboratory and stored at −20°C until analysis.

All procedures in this study fulfilled the “Reglamento de la Ley General de Salud en Materia de Investigación para la Salud” of Mexico, in particular the chapters about the ethical aspects of Research in Humans Beings, about Research in Communities, Research in minors and Research in women in fertile age and pregnant women (Diario Oficial de la Federación, febrero 1984). All methods were approved by the Ethics Committee of the Faculty of Medicine of the National Autonomous University of Mexico and research was carried out in accordance with the Declaration of Helsinki.

### Parasite detection in feces

The presence of the main intestinal parasites historically found in Xoxocotla *(Entamoeba histolytica/dispar, Entamoeba coli, Blastocystis hominis, Iodamoeba butshlii, Endolimax nana, Chilomastix mesnili, Giardia intestinalis* and *Cryptosporidium parvum*) was tested in feces by microscopy. The protists that have been associated with pathogenesis *(Entamoeba histolytica, Blastocystis hominis, Giardia intestinalis* and *Cryptosporidium parvum*) were also tested by quantitative PCR (qPCR) as previously reported (11). Briefly, samples with ~50 mg of stool were mechanically lysed using Mo Bio dry bead tubes (Mo Bio Laboratories, Inc.) in a FastPrep homogenizer (FastPrep instrument; MP Biochemicals). DNA extraction was made with the Qiagen QIAamp Fast DNA Stool Mini Kit (Qiagen) following manufacturer’s instructions. qPCR was performed on an Applied Biosystems 7500 machine using QuantiTect SYBR green master mix (Qiagen) in 10ul reaction mixture volumes with 6.25 pmol each of primers Ehd-239F–Ehd-88R, BhRDr-RD5, Giardia-80F–Giardia-127R, and CrF-CrR (Table S1). The amplification conditions consisted of 35 cycles of 1 min each at 94°C, 59°C, and 72°C, with an additional step of 95°C for 15 s, 60°C for 1 min, 95°C for 30 s, and 60°C for 15 s (55). Samples previously known to be positive for each parasite as well as standard curves using DNA from each parasite from an ATCC’s enteric protist DNA panel were included as positive controls in the qPCR plates. The difference between the average cycle threshold (CT) value of each parasite qPCR and the average CT value of the 18S rRNA gene reaction was calculated to determine the parasitic loads in each sample.

### Determination of fecal bacteria composition

DNA isolated from mothers and children fecal samples was used for the sequencing of microbial communities. For bacterial determination, samples were amplified by PCR in triplicate using bar-coded primer pairs flanking the V4 region of the 16S rRNA gene as previously described (56, 57). Each 50 μl of PCR mixture contained 22 μl of water, 25 μl of TopTaq master mix (Qiagen), 0.5 μl of each forward and reverse bar-coded primer (57), and 2 μl of template DNA. To ensure no contamination occurred, controls without template DNA were included. Amplification was performed with an initial DNA denaturation step at 95°C (5 min), 25 cycles of DNA denaturation at 95°C (1 min), an annealing step at 50°C (1 min), an elongation step at 72°C (1 min), and a final elongation step at 72°C (7 min). Amplicons displaying bands at ~250 bp on a 2% agarose gel were purified using the QIAquick PCR purification kit (Qiagen). Purified samples were quantified with PicoGreen (Invitrogen) in a Tecan M200 plate reader (excitation at 480 nm and emission at 520 nm).

For 16S rRNA gene sequencing, each PCR pool was analyzed on the Agilent Bioanalyzer using the high-sensitivity double-stranded DNA (dsDNA) assay to determine approximate library fragment size and verify library integrity. Pooled-library concentrations were determined using the TruSeq DNA sample preparation kit, version 2 (Illumina). Library pools were diluted to 4 nM and denatured into single strands using fresh 0.2 N NaOH. The final library loading concentration was 8 pM, with an additional PhiX spike-in of 20%. Sequencing was carried out using a Hi-Seq 2000 bidirectional Illumina sequencing and cluster kit, version 4 (Macrogen, Inc.).

### Determination of fecal eukaryotic composition

The composition of eukaryotic microorganisms was determined by 18S rRNA gene sequencing. DNA samples were sent to the Integrated Microbiome Resource at Dalhousie University for amplification and sequencing. The 18S rRNA gene was amplified with the primers E572F (5’ YGCGGTAATTCCAGCTC 3’) and E1009R (5’ AYGGTATCTRATCRTCTTYG 3’), and the reaction mixture included a PNA blocking primer (5’ TCTTAATCATGGCCTCAGTT 3’) to reduce amplification of mammalian sequences. Amplification was carried out in duplicate, with one reaction mixture using undiluted DNA and the other using DNA diluted 1:10 in PCR water. Amplification was conducted according to previously described protocols (58). PCR products were visualized on E-gels, quantified using Invitrogen Qubit with PicoGreen, and pooled at equal concentrations, according to a previous report (58). PhiX was spiked in at 5%, and the resulting library was sequenced at Dalhousie University on the Illumina MiSeq using the MiSeq 500-cycle reagent kit, version 2 (250 x 2).

### Bioinformatics analysis

Sequences were preprocessed, demultiplexed, denoised, quality filtered, trimmed and chimeras removed using the dada2 and vegan packages in R for 16S rRNA gene (59) or QIIME for 18S rRNA gene (60). Quality sequences were aligned to the SILVA bacterial reference alignment, and OTUs were generated using a dissimilarity cut-off of 0.03. Sequences were classified using the assignTaxonomy code and calculated alpha and beta diversities and statistic using phyloseq package in R. We estimated bacterial alpha-diversity using the Shannon index calculated from OTU relative abundances for each group. For data visualization ggplot2 package was used.

For 18S rRNA sequences, demultiplexed reads were trimmed to a uniform length of 250 bp using the FastX-Toolkit (http://hannonlab.cshl.edu/fastx_toolkit/) and clustered into OTUs using the minimum entropy decomposition (MED) method (61) as implemented in the oligotyping microbial analysis software package (62). MED performs de novo taxonomic clustering using Shannon entropy to separate biologically meaningful patterns of nucleotide diversity from sequencing noise; the processed data are partitioned into phylogenetically homogeneous units (MED nodes) for downstream bacterial diversity analyses. This analysis was carried out with the minimum substantive abundance parameter (-M) set at 250 reads. All other parameters were run with default settings; the maximum variation allowed per node - V) was automatically set at 3 nucleotides.

Representative sequences were classified by clustering against the Greengenes Database at 97% similarity (16S rRNA gene [63]) or SILVA release 123 at 99% similarity (18S rRNA gene [64]). The 16S rRNA gene data set was filtered to remove mitochondrion and chloroplast sequences and OTUs present in fewer than three samples. The 18S rRNA gene data set was filtered to remove mammalian and plant sequences and all OTUs present in fewer than three samples. Both data sets were filtered to exclude singletons and doubletons.

### Statistical analysis

Differences in frequencies for categorical and continuous variables between cases and controls were evaluated using the chi-squared and Student’s t test, respectively.

Microbial diversity and the relative abundances of bacterial and eukaryotic taxa was evaluated using phyloseq (65), along with additional R-based computational tools (66–72). PCoAs were conducted using phyloseq (Bray-Curtis dissimilarities as distance metric) on both variance-stabilizing-transformed and rarefied OTU matrices and then statistically confirmed by a permutational multivariate analysis of variance (PERMANOVA) to confirm that our results were not a consequence of heteroscedastic dispersion between groups (65). The Shannon and Chao1 alpha diversity indexes were calculated using phyloseq and statistically confirmed by the Mann-Whitney test (GraphPad Prism software, version 5c).

The R packages DESeq2 (72) and MaAsLin (73) were used to calculate differentially abundant OTUs. Correlation analysis was performed using the bicor method in the R package microbiome to correlate the 100 most abundant OTUs from the 16S and 18S rRNA gene data sets. Features in the analysis were included as OTUs and as OTUs combined into taxonomic families.

## ACKNOWLEDGEMENTS

This work was funded by grant numbers IN226511 and IN218214 from PAPIIT program at UNAM: grant FIS/IMSS/PROT/1368 from IMSS, and grants numbers 140990, 272601, 283522, and 257091 from the National Council of Sciences and Technology in Mexico (CONACyT) to C.X.-G. and J.T., Canadian Institutes for Health Research grants to B.B.F. and a Human Frontier Science Program grant to L.W.P. (grant number RGY0078/2015). The funders had no role in study design, data collection and interpretation, or the decision to submit the work for publication. O.P.-R. (proposal number 208253) and M.E.N.-R. (proposal number 235618) each received a 1-year scholarship from the Estancias Posdoctorales al Extranjero para la Consolidacion de Grupos de Investigación program of CONACyT. We thank all study participants from Xoxocotla as well as the personnel involved in this cohort.

## Author contributions

O.P.-R., M.N.-R., M.C.A., E.B., C.X., and B.B.F. designed the study. A.V.-S., P.M., J.T, O.P.-R., M.N.-R., E.G., E. R, U.M, L.R.-V., E.H, and C.X. coordinated and facilitated the cohort study in Xoxocotla. P.M. and C.X. conducted medical examinations and Rome III questionnaires. A.V.-S. curated the database and metadata. M.C.A. and L.W.P. optimized sequencing strategies. O.P.-R. and M.N.-R. prepared samples for sequencing analysis. M.C.A. and E.M. performed the bioinformatics analysis of sequencing data. M.C.A. and I.L.-L. designed and performed statistical analyses and created figures for the paper. O.P.-R., I. L.-L. and M.C.A. wrote the paper. M.E.N.-R., I.L.-L., L.W.P, L.R., C.X.-G., M.C.A. and B.B.F. edited.

